# Leveraging inter-individual transcriptional correlation structure to infer discrete signaling mechanisms across metabolic tissues

**DOI:** 10.1101/2023.05.10.540142

**Authors:** Mingqi Zhou, Ian J. Tamburini, Cassandra Van, Jeffrey Molendijk, Christy M Nguyen, Ivan Yao-Yi Chang, Casey Johnson, Leandro M. Velez, Youngseo Cheon, Reichelle X. Yeo, Hosung Bae, Johnny Le, Natalie Larson, Ron Pulido, Carlos Filho, Cholsoon Jang, Ivan Marazzi, Jamie N. Justice, Nicholas Pannunzio, Andrea Hevener, Lauren M. Sparks, Erin E. Kershaw, Dequina Nicholas, Benjamin Parker, Selma Masri, Marcus Seldin

## Abstract

Inter-organ communication is a vital process to maintain physiologic homeostasis, and its dysregulation contributes to many human diseases. Beginning with the discovery of insulin over a century ago, characterization of molecules responsible for signal between tissues has required careful and elegant experimentation where these observations have been integral to deciphering physiology and disease. Given that circulating bioactive factors are stable in serum, occur naturally, and are easily assayed from blood, they present obvious focal molecules for therapeutic intervention and biomarker development. For example, physiologic dissection of the actions of soluble proteins such as proprotein convertase subtilisin/kexin type 9 (*PCSK9*) and glucagon-like peptide 1 (*GLP1*) have yielded among the most promising therapeutics to treat cardiovascular disease and obesity, respectively^1–4^. A major obstacle in the characterization of such soluble factors is that defining their tissues and pathways of action requires extensive experimental testing in cells and animal models. Recently, studies have shown that secreted proteins mediating inter-tissue signaling could be identified by “brute-force” surveys of all genes within RNA-sequencing measures across tissues within a population^5–9^. Expanding on this intuition, we reasoned that parallel strategies could be used to understand how individual genes mediate signaling across metabolic tissues through correlative analyses of gene variation between individuals. Thus, comparison of quantitative levels of gene expression relationships between organs in a population could aid in understanding cross-organ signaling. Here, we surveyed gene-gene correlation structure across 18 metabolic tissues in 310 human individuals and 7 tissues in 103 diverse strains of mice fed a normal chow or HFHS diet. Variation of genes such as *FGF21, ADIPOQ, GCG* and *IL6* showed enrichments which recapitulate experimental observations. Further, similar analyses were applied to explore both within-tissue signaling mechanisms (liver *PCSK9*) as well as genes encoding enzymes producing metabolites (adipose *PNPLA2*), where inter-individual correlation structure aligned with known roles for these critical metabolic pathways. Examination of sex hormone receptor correlations in mice highlighted the difference of tissue-specific variation in relationships with metabolic traits. We refer to this resource as **<u>G</u>**ene-**<u>D</u>**erived **<u>C</u>**orrelations **<u>A</u>**cross **T**issues (GD-CAT) where all tools and data are built into a web portal enabling users to perform these analyses without a single line of code (gdcat.org). This resource enables querying of any gene in any tissue to find correlated patterns of genes, cell types, pathways and network architectures across metabolic organs.

## Results

### Construction of a web tool to survey transcript correlations across tissues and individuals (GD-CAT)

Previous studies have established that “brute force” analyses of correlation structure across tissues from population expression data can identify new several known and mechanisms of organ vcross-talk. These were accomplished by surveying the global correlation structure using all genes, whereby skewed upper-limits of significance distributions were sufficient to prioritize proteins which elicit signaling^5–9^. Following this intuition, we hypothesized that a paralleled but alternative approach to inter-individual correlation structure could be exploited to understand the functional consequences of specific genes. Our initial goal was to establish a user-friendly interface where all of these analyses and gene-centric queries could be performed without running any code. To accomplish this, we assembled a complete analysis pipeline (Fig 1A) as a shiny-app and docker image hosted in a freely-available web address (gdcat.org). Here, users can readily-search gene correlation structure between individuals from filtered human (gene-by-tissue expression project - GTEx) and mouse (hybrid mouse diversity panel - HMDP) across tissues. GTEx is presently the most comprehensive pan-tissue dataset in humans^10^, which was filtered for individuals where most metabolic tissues were sequenced^9^. Collectively, this dataset contains 310 individuals, consisting of 210 male and 100 female (self-reported) subjects between the ages of 20-79. Data from the HMDP consisted of 96 diverse mouse strains fed a normal chow (5 tissues) or high-fat/high-sucrose diet (7 tissues) as well as carefully characterized clinical traits ^11–16^. Initially, users select a given species, followed by reported sex or diet (mouse) which loads the specified environment. Subsequent downstream analyses are then implemented accordingly from a specific gene in a given tissue.. This selection prompts individual gene correlations across all other gene-tissue combinations using biweight midcorrelation^17^. From these charts, users are able to select a given tissue, where gene set enrichment analysis testing using clusterprofiler^18^ and enrichR ^19^ are applied to the correlated set of genes to determine the positively (activated) and negatively (suppressed) pathways which occur in each tissue. In addition to general queries of gene ∼ gene correlation structure, comparison of expression changes are also visualized between age groups as well as reported sexes. In addition, we included the top cell-type abundance correlations with each gene. To compute cell abundance estimates from the same individuals, we used single-nucleus RNA-seq available from GTEx^20^ and applied cellular deconvolution methods to the bulk RNA-seq^21^ (methods). Comparison of deconvolution methods^21^ showed that DeconRNASeq^22^ captured the most cell types within several tissues (Supplemental Figures 1-3) and therefore was applied to all tissues where sn-RNA-seq was available. We note that visceral adipose, subcutaneous adipose, aortic artery, coronary artery, transverse colon, sigmoid colon, the heart left ventricle, the kidney cortex, liver, lung, skeletal muscle, spleen, and small intestine are the only tissues where sn-seq is available and not other tissues, such as brain, stomach and thyroid.

**Figure 1.**
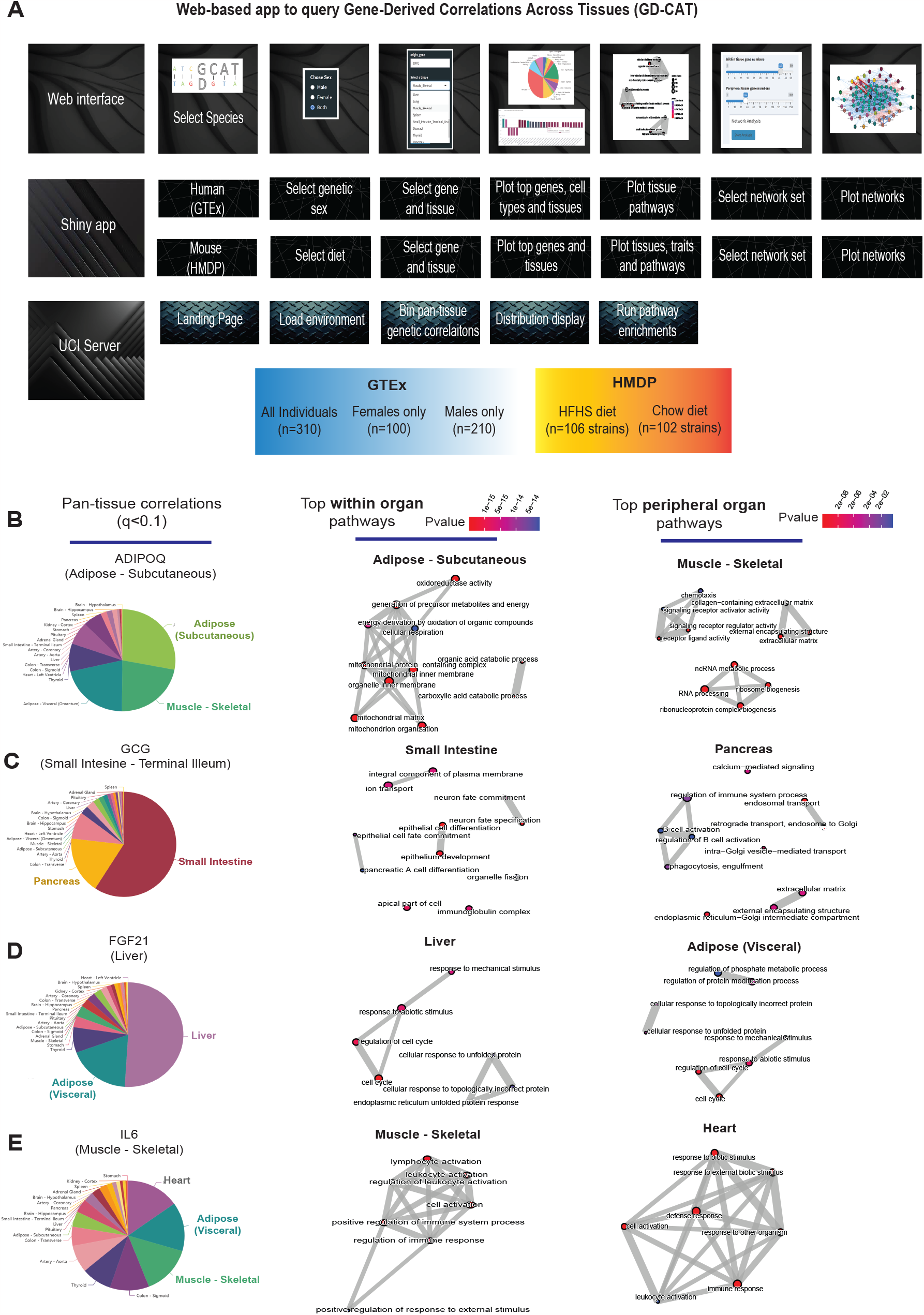
Web tool overview and inter-individual correlation structure of established endocrine proteins. A, Web server structure for user-defined interactions, as well as server and shiny app implementation scheme for GD-CAT. B, All genes across the 18 metabolic tissues in 310 individuals were correlated with expression of *ADIPOQ* in subcutaneous adipose tissue, where a qvalue cutoff of q<0.1 showed the strongest enrichments with subcutaneous and muscle gene expression (pie chart, left). Gene set enrichment analysis (GSEA) was performed using the bicor coefficient of all genes to *ADIPOQ* using gene ontology biological process annotations and network construction of top pathways using clusterprofiler, where pathways related to fatty acid oxidation were observed in adipose (left) and chemotaxis/ECM remodeling in skeletal muscle (right). B-D, The same qvalue binning, top within-tissue and top peripheral enrichments were applied to intestinal *GCG* (B), liver *FGF21* (C) and muscle *IL6* (D). For these analyses all 310 individuals (across both sexes) were used and qvalue adjustments calculated using a Benjamini-Hochberg FDR adjustment.

We initially examined pan-tissue transcript correlation structures for several well-established mechanisms of tissue crosstalk via secreted proteins which contribute to metabolic homeostasis. Here, binning of the significant tissues and pathways related to each of these established secreted proteins resembled their known mechanisms of action (Fig 1B-E). For example, variation with subcutaneous adipose expression of *ADIPOQ* was enriched with genes in several metabolic tissues where it has been known to act (Fig 1B, left). In particular, subcutaneous adipose *ADIPOQ* expression correlated with fatty acid oxidative process within adipose (Fig 1B, middle) and was enriched with ECM, chemotaxis and ribosomal biogenesis in skeletal muscle (Fig 1B, right). These correlated pathways align with the established physiologic roles of the protein in that fat secreted adiponectin when oxidation is stimulated^23,24^ and muscle is a major site of action^25^. Beyond adiponectin, inter-individual correlation structure additionally recapitulated broad signaling mechanisms for other relevant endocrine proteins. For example, intestinal *GCG* (encoding GLP1, Fig 1C), liver *FGF21* (Fig 1D) and skeletal muscle *IL6* (Fig 1E) showed binning patterns and pathway enrichments related to their known functions in pancreas^1,26^, adipose tissue^27,28^ and other metabolic organs^29^, respectively. These analyses and web tool show some examples of exploring transcriptional correlation structure to confirm and identify mechanisms of signaling, where we note that additional limitations should be considered.

### Pathway-based examination of gene correlation structure and significance thresholds across tissues

While the select observations shown in Fig1 provide examples of support in exploring correlation structure of genes across interindividual differences to investigate endocrinology, several limitations in these analyses should be considered. First, an additional explanation for a given gene showing strong correlation between the tissues could arise from a general pattern of correlation between the two tissues and not necessarily due to the discrete signaling mechanisms. In previous studies surveying correlation structure and network model architectures in the HMDP and STARNET populations, genes appeared generally stronger correlated between liver and adipose tissue compared to all other organ combinations explored^5–7^. To investigate this global pattern of gene correlation structure between metabolic organs, we selected key GO terms, KEGG pathways and randomly sampled equal numbers of genes and evaluated relative significance of inter-tissue correlations across multiple statistical thresholds. These analyses suggested that usage of empirical student correlation pvalues recapitulated a clear pattern of inter-tissue correlations between pathways (Fig 2). For example, comparison of the number of genes achieving significance of correlation between tissues among select GO terms revealed that tissues such as adipose and muscle appeared more correlated than spleen and other tissues at pvalues less than 1e-3 (Fig 2A, left column). These global patterns of gene correlation between tissues among select pathways were reduced when the pvalue threshold was lowered to 1e-6 (Fig 2A, middle column) or qvalue adjustments (methods) were performed (Fig 2A, right two columns). For these reasons, only qvalue adjusted value were used and implemented into pie charts providing the tissue-specific occurrences of correlated genes at 3 thresholds (q<0.1, q<0.01, q<0.001) within the web tool. Next, in order to further evaluate these global patterns of innate transcript correlation structure and determine whether they reflected concordance between known metabolic pathways or innate to the dataset used, tissues were rank-ordered by the number of genes which meet pvalue thresholds and compared to randomly sampled genes of similar pathway sized (Fig 2B). Among KEGG Pathways selects (hsa04062 − Chemokine signaling pathway, hsa04640 − Hematopoietic cell lineage and hsa00190 − Oxidative phosphorylation), the top-ranked organs by correlated gene numbers differed (Skeletal muscle, Colon and Thyroid, respectively); however, a general trend of specific tissues ranking higher than others were observed (Fig 2B). For example, skeletal muscle and heart appeared among the strongest correlated across pathways and organs, compared to kidney cortex and spleen which were observed to rank among the lowest (Fig 2B, pathways). We note that when the same analysis was performed on randomly sampled genes from each organ consisting of the same number as genes within each KEGG pathway, these rankings and number of significant correlating genes were no longer observed (Fig 2B, random genes), suggesting that in certain instances differences between organs in general connectivity to others might reflect concordance between known pathways. It is important to consider here that for the organs ranking lower, the lack of relative correlating numbers is likely due to sparsity of available data and not necessarily general patterns of gene correlation. This point is supported by the fact that among the lowest-ranked 33% of tissues across pathways, we observed a significant negative overall correlation (bicor = -0.45, pvalue = 2.3e-5) between number of NA values per individual and the gene count for significance shown in Fig 2B. This negative correlation between missing data and number of significant correlations for pathways across tissues was not observed when binning the top 33% (bicor = 0.09, pvalue = 0.42) or middle 33% (bicor = -0.12, pvalue = 0.27) of organs. Collectively, these analyses show that innate correlations structures exist between organs which differ depending on pathways investigated and that tissues which don’t show broad correlation structure could potentially be attributed to areas of missing data among GTEx.

**Figure 2.**
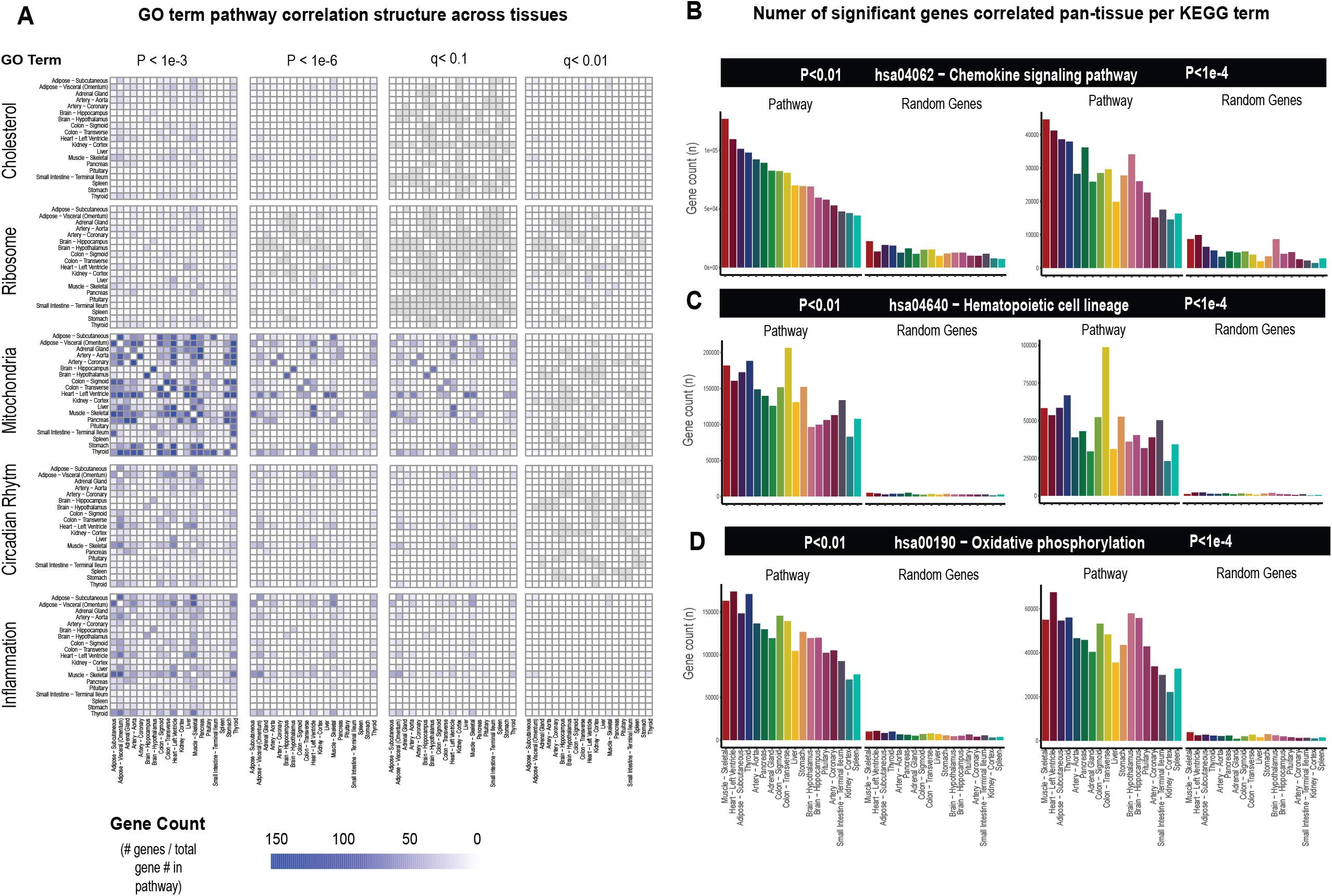
Tissue-specific contributions to pan-organ gene-gene correlation structure. A Heatmap showing all the number of gene-gene correlations across tissues which achieve significance relative to total number of genes in each pathway at biweigth midcorrelation student pvalue < 1e-3 (left column), pvalue < 1e-6 (left middle column) of BH-corrected qvalue <0.1 (right middle column) or BH-corrected qvalue<0.01 (right column). Within-tissue correlations are omitted from this analysis. B-D, Genes corresponding to each KEGG pathway shown were correlated both within and across all other organs where the number of genes which meet each students pvalue threshold are shown (y-axis). Tissues (x-axis) are rank-ordered by the number of genes which correlate for hsa04062 − Chemokine signaling pathway at pvalue<0.01 and shown for other KEGG terms, hsa04640 − Hematopoietic cell lineage (C) and hsa00190 − Oxidative phosphorylation (D) and additionally pvalue<1e-4 (right side).

**Figure 3.**
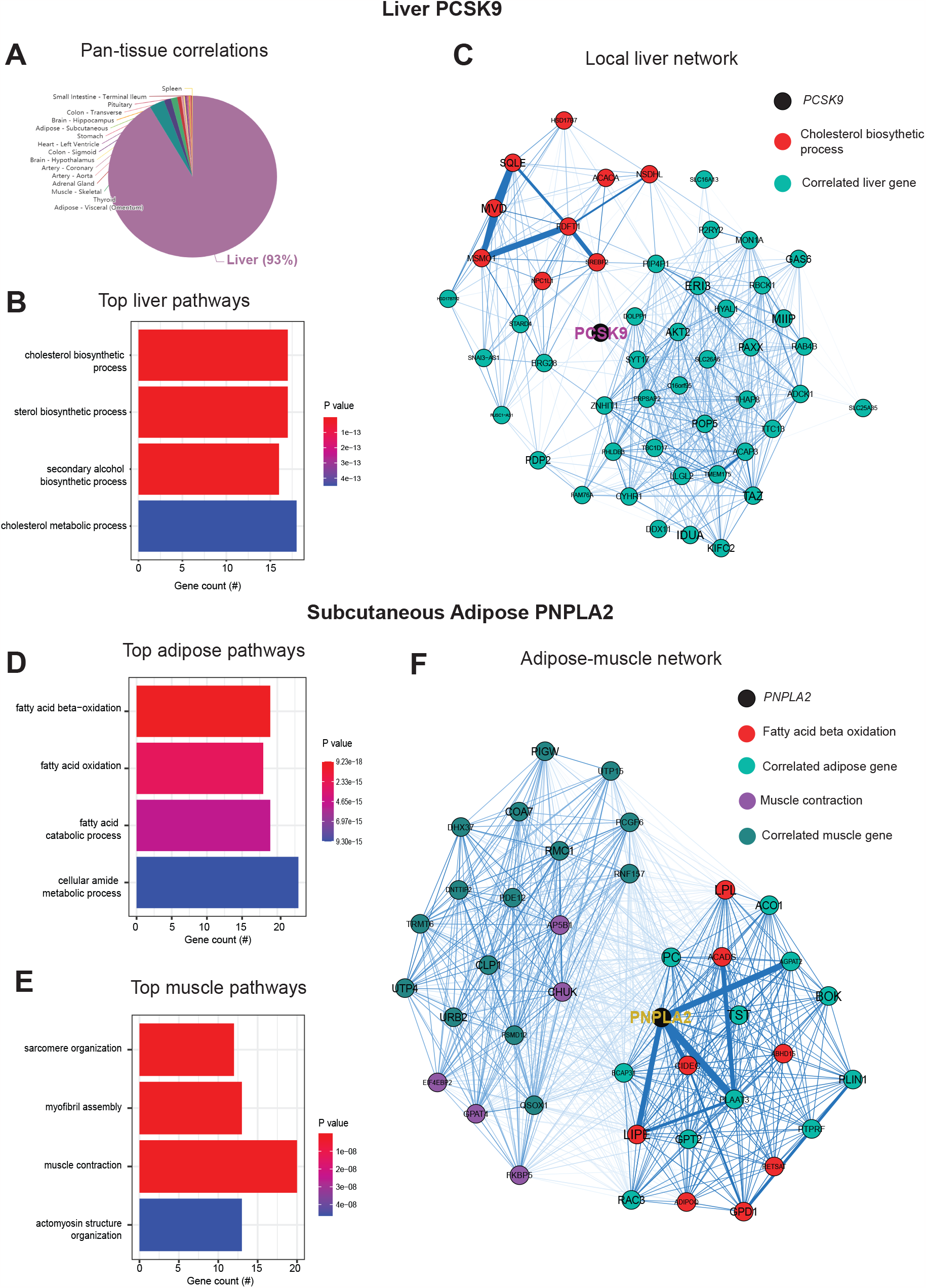
Inter-individual transcript correlation structure and network architecture of liver *PCSK9* and adipose *PNPLA2*. A, distribution of pan-tissue genes correlated with liver *PCSK9* expression (q<0.1), where 93% of genes were within liver (purple). B, Gene ontology (BP) overrepresentation test for the top 500 hepatic genes correlated with *PCSK9* expression in liver. C, Undirected network constructed from liver genes (aqua) correlated with *PCSK9*, where those annotated for “cholesterol biosynthetic process” are colored in red. D-E, over-representation tests corresponding to the top-correlated genes with adipose (subcutaneous) *PNPLA2* expression residing in adipose (D) or peripherally in skeletal muscle (E). F, Undirected network constructed from from the strongest correlated subcutaneous adipose tissue (light aqua) and muscle genes (dark blue) with PNPLA2 (black), where genes corresponding to GO terms annotated as “fatty acid beta oxidation” or “Muscle contraction” are colored purple or red, respectively. For these analyses all 310 individuals (across both sexes) were used and qvalue adjustments calculated using a Benjamini-Hochberg FDR adjustment. Network graphs generated based in Biweight midcorrelation coefficients, where edges are colored blue for positive correlations or red for negative correlations. Network edges represent positive (blue) and negative (red) correlations and the thicknesses are determined by coefficients. They are set for a range of bicor=0.6 (minimum to include) to bicor=0.99

### PSCK9 signaling and lipid exchange between adipose and muscle apparent in simple network models of correlation structure

Next, we wanted to ask whether our approach of analyzing inter-individual correlation structure across tissue for endocrine proteins was also sufficient to define within-tissue signaling mechanisms or actions of enzymes producing metabolites that signal across organs. Dissimilar to the cross-tissue distributions of significance in Fig 1, the same analysis of liver *PCSK9* highlighted exclusively liver genes which were varied together (Fig 2A), in particular those involved in cholesterol metabolism/homeostasis (Fig 2B). Consistent with the established role for PCSK9 as a primary degradation mechanism of LDLR^4,30^, network model construction of correlated genes highlighted the gene as a central node linking cholesterol biosynthetic pathways with those involved in other metabolic pathways such as insulin signaling (Fig 2C). Given that organ signaling via metabolites comprises many critical processes among multicellular organisms, our next goal was to apply this gene-centric analyses to established mechanisms of metabolite signaling. The gene *PNPLA2* encodes adipose triglyceride lipase (ATGL) which localizes to lipid droplets and breaks down triglycerides for oxidation or mobilization as free fatty acids for peripheral tissues^31^. Variation in expression of *PNPLA2* showed highly significant enrichments with beta oxidation pathways in adipose tissue (Fig 2D). Muscle pathways enriched for the gene were represented by sarcomere organization and muscle contraction (Fig 2F). Construction of an undirected network from these expression data placed the gene as a central node between the two tissues, linking regulators of adipose oxidation (Fig 2F, red) to muscle contractile process (Fig 2F, purple) where additional strongly co-correlated genes were implicated as additional candidates (Fig 2F). In sum, these analyses provide two examples of within-liver signaling via *PCSK9* and adipose-muscle communication through *PNPLA2* where the top-correlated genes and network models recapitulate known mechanisms. Given the utility of these undirected network models, a function in GD-CAT was added to enable users to generate network models for any gene-tissue combination and select parameters such as number of within-tissue and peripheral correlated genes to include.

### Inter-individual correlation analysis of hybrid mouse diversity panel highlights tissue- and diet-specific phenotype relationships with sex hormones

Genetic reference panels in model organisms, such as mice, present appeal in studying complex traits in that environmental conditions can be tightly controlled, tissues and invasive traits readily accessible and the same, often renewable, genetic background can be studied and compared among multiple exposures such as diets or drug treatments ^15,32–34^. For this resource, we utilized data from the HMDP fed a normal chow^15,16^ or HFHS diet for 8 weeks^11–14^. While the number of tissues available was less than in GTEx, these panels allow for comparison of how gene correlations shift depending on diet. Therefore, queries of gene correlation queries in mice were segregated into either chow or HFHS diet and an additional panel to download a table or visualize the relationship between genes and clinical measures was added. The inferred abundances of cell types from each individual are correlated across user-defined genes, with the bicor coefficient plotted for each cell type.

One advantage of hybrid mouse diversity panel data compared to GTEx is the abundance of phenotypic measures available within each cohort. To show the utility of examining correlations within this reference panel, we selected sex hormone receptors androgen receptor (*Ar*), estrogen receptor alpha (*Esr1*) or estrogen receptor beta (*Esr2*) and binned the top 10 phenotypes which were correlated. These analyses were segregated based on where sex hormones were expressed (either liver or adipose tissue) or dietary regiment of the ∼100 strains (normal chow or HFHS diet). This analysis demonstrated the difference in relationships between tissue location of sex hormone receptor and dietary context with metabolic traits. For example, expression of *Ar* in adipose tissue among HMDP mice fed a HFHS diet was negatively correlated with fat mass and body weight traits, whereas expression in liver oppositely correlated with the same traits in a positive direction (Fig 4A). The top traits which correlated also differed by tissue or expression for *Ar*, such as plasma lipid parameters in adipose tissue compared to blood cell traits in chow-fed mice (Fig 4A). We note that among the three hormone receptors investigated, *Esr2* appeared the most consistently correlated between tissues and diets with metabolic traits (Fig 4B). Expression of *Esr1* also showed a clear tissue and diet difference in the traits which were the most strongly co-regulated. Under HFHS dietary conditions, a negative correlation with insulin and fat pad weights were observed exclusively with adipose expression, while positive correlations with liver lipids were observed with expression in liver (Fig 4C). These analyses highlight how phenotype correlations in mouse populations can help to determine contexts relevant for gene regulation and point to the diversity of potential contexts relevant for sex hormone receptors in metabolic tissues.

**Figure 4.**
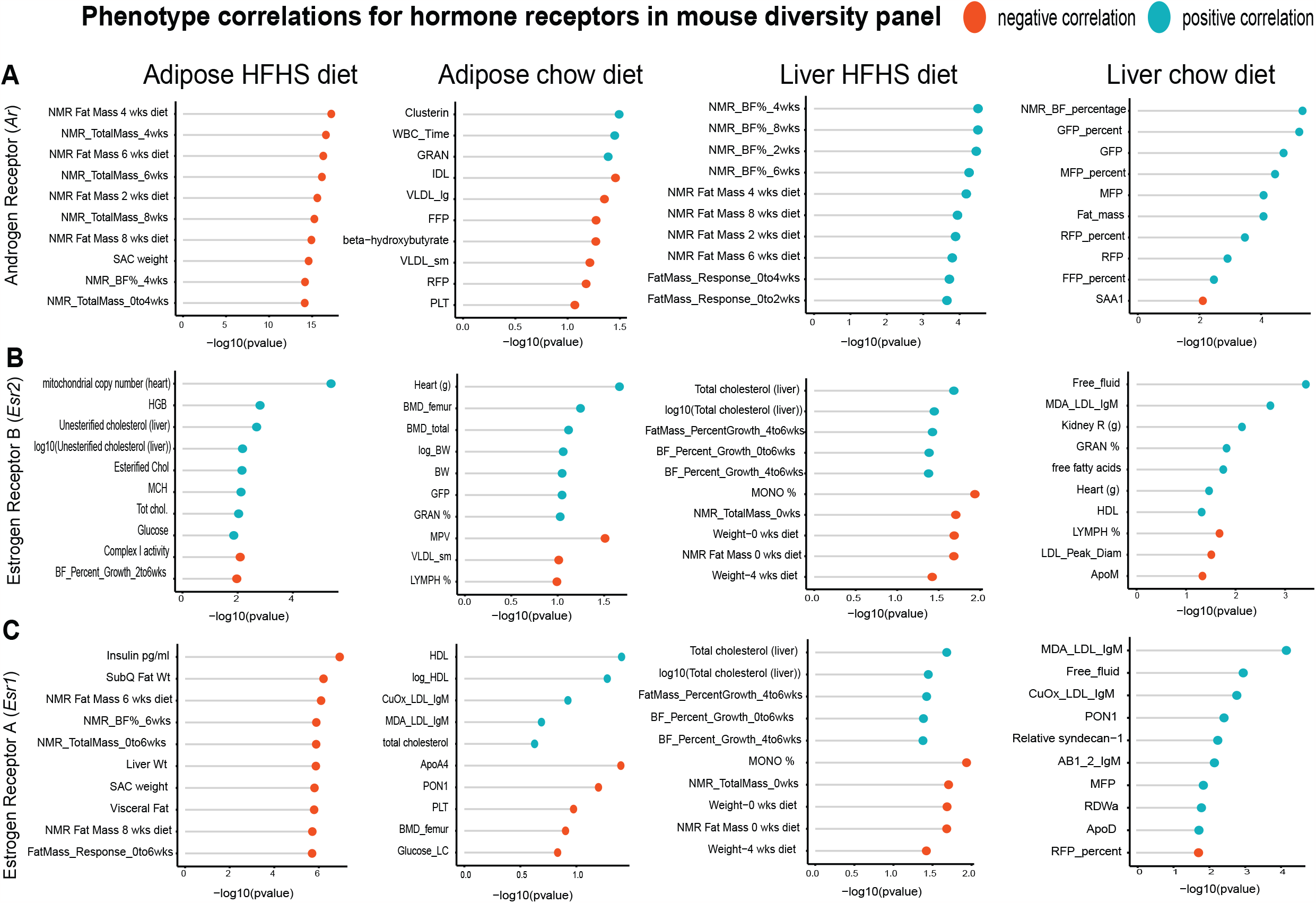
HMDP tissue- and diet-specific correlations of sex hormone receptors. The top 10 phenotypic traits which correlated to expression of androgen receptor (A), estrogen receptor 1 (B) or estrogen receptor 2 (C) colored by direction in the hybrid mouse diversity panel. Positive correlations are shown in light blue and negative correlations as sunset orange, where phenotypes (y-axis) are ordered by significance (x-axis, -log(pvalue) of correlation). Correlations are segregated by whether sex hormone receptors are expressed by gonadal adipose tissue (left two columns) in ∼100 HMDP strains fed a HFHS diet (left), normal chow diet (left middle) or liver-expressed receptors fed a HFHS diet (right middle) or normal chow diet (right).

## Discussion

### Limitations and Conclusions

Here, we provide a new resource to explore correlations across organ gene expression in the context of interindividual differences. We highlight areas where these align with established and relevant mechanisms of physiology and suggest that similar explorations could be used as a discovery tool. Several key limitations should be considered when exploring GD-CAT for mechanisms of inter-tissue signaling though. Primarily, the fact that correlation-based analyses could reflect both causal or reactive patterns of variation. While several statistical methods such as mediation^35,36^ and mendelian randomization^37,38^ exist to further refine causal inferences, likely the only definitive method to distinguish is in carefully-designed experimentation. Further, analyses of genetic correlation (ex. correlations considering genetic loci to infer causality) also present appeal in refining some causal mechanisms. Correlation between molecular and phenotypic variables can occur for a variety of reasons, not just between their individual relationships, but often more broadly, from a variety of complex genetic and environmental factors. Further, many correlations tend to be dominated by genes expressed within the same organ. This could be due to the fact that, within-tissue correlations could capture both the pathways regulating expression of a gene, as well as potential consequences of changes in expression/function, and distinguishing between the two presents a significant challenge. For example, a GD-CAT query of insulin (*INS*) expression in pancreas shows exclusive enrichments in pancreas and corresponding pathway terms reflect regulatory mechanisms such as secretion and ion transport (Supplemental Fig 4). Representation of given genes may also differ significantly depending on the dataset used. For example, while queries of other tissues for the critical X Inactive Specific Transcript (*XIST*), in liver no significant correlations appear. This is due to the fact that the gene operates in a sex-dependent manner, where females are significantly less represented in GTEx and liver exists as a sparser tissue compared to others (Fig 2). In addition, the analyses presented are derived from differences in gene expression across individuals which arise from complex interaction of genetic and environmental variables. Expression of a gene and its corresponding protein can show substantial discordances depending on the dataset used. These have been discussed in detail^39–41^, but ranges of co-correlation can vary widely depending on the datasets used and approaches taken. We note that for genes encoding proteins where actions from acute secretion grossly outweigh patterns of gene expression, such as insulin, caution should be taken when interpreting results. As the depth and availability of tissue-specific proteomic levels across diverse individuals continues to increase, an exciting opportunity is presented to explore the applicability of these analyses and identify areas when gene expression is not a sufficient measure. For example, mass-spec proteomics was recently performed on GTEx^42^; however, given that these data represent 6 individuals, analyses utilizing well-powered inter-individual correlations such as ours which contain 310 individuals remain limited in applications.

The queries provided in GD-CAT use fairly simple linear models to infer organ-organ signaling; however, more sophisticated methods can also be applied in an informative fashion. For example, Koplev et al generated co-expression modules from 9 tissues in the STARNET dataset, where construction of a massive Bayesian network uncovered interactions between correlated modules^6^. These approaches expanded on analysis of STAGE data to construct network models using WGCNA across tissues and relating these resulting eigenvectors to outcomes^43^. The generalized approach of constructing cross-tissue gene regulatory modules presents appeal in that genes are able to be viewed in the context of a network with respect to all other gene-tissue combinations. In searching through these types of expanded networks, individuals can identify where the most compelling global relationships occur. One challenge with this type of approach; however, is that coregulated pathways and module members are highly subjective to parameters used to construct GRNs (for example reassignment threshold in WGCNA) and can be difficult in arriving at a “ground truth” for parameter selection. We note that the WGCNA package is also implemented in these analyses, but solely to perform gene-focused correlations using biweight midcorrelation to limit outlier inflation. While the midweight bicorrelation approach to calculate correlations could also be replaced with more sophisticated models, one consideration would be a concern of overfitting models and thus, biasing outcomes.

In another notable example MultiCens was developed as a tool to uncover communication between genes and tissues and applied to suggest central processes which exist in multi-layered data relevant for Alzheimer’s disease^44^. In addition, Jadhav and colleagues adopted a machine learning approach to mine published literature for relationships between hormones and genes^45^. Further, association mapping of plasma proteomics data has been extensively applied and intersection with genome-wide association disease loci has offered intriguing potential disease mechanisms^46,47^. Another common application to single-cell sequencing data is to search for overrepresentation of known ligand-receptor pairs between cell types^48^. These and additional applications to explore tissue communication/coordination present unique strengths and caveats, depending on the specific usage desired. Regardless of methods used to decipher, one important limitation to consider in all these analyses is the nature of underlying data. For example, our evaluation of GTEx data structure suggested that important organs such as spleen and kidney were insufficient due to availability in matching expression data between individuals. Further, GTEx sample vary as to the collection times, sample processing times and other important parameters such as cause of death. Mouse population data such as the HMDP or BxD cohorts offer appeal in these regards, as environmental conditions and collection times are easily fixed. Regardless, careful consideration of how data was generated and normalized are fundamental to interpreting results.

In sum we demonstrate that adopting a gene-centric approach to surveying correlation structure of transcripts across organs and individuals can inform mechanism of coordination between metabolic tissues. Initially, we queried several well-established and key mediators of physiologic homeostasis, such as *FGF21, GCG* and *PCSK9*. These approaches are further suggested to be applicable to mechanisms of metabolite signaling, as evident by pan-tissue investigation of adipose *PNPLA2*. Exploration of hybrid mouse diversity panel data highlighted the diverse phenotype correlations depending on tissue and diet for sex hormone receptors. To facilitate widespread access and use of this transcript isoform-centric analysis of inter-individual correlations, a full suite of analyses such as those performed here can be performed from a lab-hosted server (gdcat.org) or in isolation from a shiny app or docker image.

## Material and methods

### Availability of web tool and analyses

All analyses, datasets and scripts used to generate the associated web tool (GD-CAT) can be accessed via: https://github.com/mingqizh/GD-CAT or within the associated docker image. In addition, access to the GD-CAT web tool is also available through the web portal gdcat.org. This portal was created to provide a user-friendly interface for accessing and using the GD-CAT tool without the need to download or install any software or packages. Users can simply visit the website, process data and start using the tool.

Corresponding tutorial and the other resources were made available to facilitate the utilization of the web tool on GitHub. The interface and server of the web were built and linked based on the shiny package using R (v. 4.2.0). Shiny package provides a powerful tool for building interactive web applications using R, allowing for fast and flexible development of custom applications with minimal coding required.

### Pathway-specific gene correlations across tissues

Detailed scripts and analyses for pathway-specific investigations across tissues in Fig2 are provided in: https://github.com/itamburi/gtex-app-kegg-pathways. Briefly, to interrogate broad tissue correlation structure, the number of genes which passed each biweight midcorrelation pvalue cutoff are shown normalized to the total number of genes corresponding to that pathway term. Pathways were selected by accessing all available GO annotations for all genes using the Universal Protein Resource^49^ and subletting genes where a given term is listed. To determine which tissues show the most co-correlation across genes and organs, KEGG terms shown were selected and each corresponding gene-tissue combinations were correlated. Tissues were then binned by the number of significant correlations which occurring both within and across organs among each selected KEGG pathway at indication correlation pvalues. Rank-ordering on the figure was shown by chemokine signaling at P<0.01 and each term was compared to a randomly sampled set of genes corresponding to the same number contained in each pathway.

### Data sources and availability

All human data used in this study can be immediately accessed via web tool or docker to facilitate analysis. Metabolic tissue data was accessed through GTEx V8 downloads portal on August 18, 2021 and previously described^9,10^. These raw data can also be readily accessed from the associated R-based walkthrough: https://github.com/Leandromvelez/myokine-signaling. Briefly, these data were filtered to retain genes which were detected across tissues where individuals were required to show counts > 0 across all data. Given that our goal was to look across tissues at enrichments, this was done to limit spurious influence of genes only expressed in specific tissues in specific individuals.

Hybrid mouse diversity panel data was collected from previously described studies^11,15,16,34^ and inter-individual differences were compared at the strain-level to maximize possible comparisons between historical data.

### Correlation analyses across tissues

biweight midcorrelation coefficients and corresponding p-values within and across tissues were generated using WGCNA bicorandpvalue() function^17^. We note that while the WGCNA package was used to calculate coefficients and corresponding students pvalues, this generalized framework does not utilize any module generation. Associated qvalue adjustments were applied using the Benjamini-Hochberg FDR from the R package “stats”. These BH adjustments, as opposed to standard qvalue adjustments, were selected given their efficiency in CPU usage on the hosted server.

### Pathway enrichment analyses

Pathway enrichments were generated using gene set enrichment analyses available from the r package clusterprofiler. Specifically, the bicor coefficients were used as the rank-weight of each gene and enrichment tests performed by permuting against the human or mouse reference transcriptome. Terms used for the enrichment analyses were derived from Gene Ontology (Biological Process, Cellular Component and Molecular Function) which were accessed using the R package enrichR. For this analysis and on the available app, input genes were determined at indicated qvalue threshold.

### Deconvolution of bulk tissue seq data on web tool

All scripts and deconvolution data produced is available at: https://github.com/cvan859/deconvolution. Briefly, sn-RNA-seq data was accessed from the Human cell atlas^20^ for matching organ datasets with metabolic tissues. From these data, 4 deconvolution methods were applied using ADAPTS^21^ where DeconRNA-Seq^22^ was selected for its ability to capture the abundances of the most cell types across tissues such as liver heart and skeletal muscle (Supplemental Fig 1-3). The full combined matrix was assembled for DeconRNA-Seq results across individuals in GTEx where correlations between cell types and genes was performed also using the bicorandpvalue() in WGCNA^17^.

## Acknowledgements

We acknowledge the following funding sources for supporting these studies: MZ, CF, IJT, CMN, LMV, CV, CJ and MMS were supported by NIH grants HL138193, DK130640 and DK097771. ALH is supported by NIH grants U54 DK120342, R01 DK109724, and P30 DK063491. NRP is supported by NIH grant R37 CA266042. HB was supported by National Research Foundation of Korea (2021R1A6A3A14039132). JL was supported by an NIH grant F31DK134173-01A1. CJ was supported by AASLD Foundation Pinnacle Research Award in Liver Disease, Edward Mallinckrodt, Jr. Foundation Award, and an NIH grant AA029124.

## Author contributions

MZ, IJT, CV and MMS accessed raw data, performed analyses and drafted the manuscript. MZ assembled the shiny application, where IV and RP designed the UCI infrastructure to enable access. CJ, CMN, LMV, JM, CF, IC, RY, HB, JL, NL, IM, AH, LMS, JNJ, EEK, IM, NP, DN and BM provided critical insight into data use and interpretation, as well as guided the study. CJ, NP and SM guided tool design involving metabolite signaling and circadian rhythms, respectively, as well as provided app infrastructure. All authors have approved the current manuscript. All authors read and approved this manuscript.

## Conflict of interest

The authors have no conflicts of interest to declare

## Figure Legends

**Supplemental Figure 1:** Performance across 4 methods of cell-type deconvolution where relative proportions of cells (y-axis) are shown for all cell types annotated in single-cell reference (x-axis) in Liver.

**Supplemental Figure 2:** Performance across 4 methods of cell-type deconvolution where relative proportions of cells (y-axis) are shown for all cell types annotated in single-cell reference (x-axis) in Heart.

**Supplemental Figure 3:** Performance across 4 methods of cell-type deconvolution where relative proportions of cells (y-axis) are shown for all cell types annotated in single-cell reference (x-axis) in Skeletal Muscle.

**Supplemental Figure 4:** Pancreatic *INS* expression correlations across tissues in GTEx were binned according to q<0.1 (top) and corresponding pancreatic GSEA network graph is shown (bottom)

## Notes

### Competing Interest Statement

The authors have declared no competing interest.

### Summary of Updates

resubmitted

